# RumimiR: a detailed microRNA database focused on ruminant species

**DOI:** 10.1101/561720

**Authors:** Bourdon Céline, Bardou Philippe, Aujean Etienne, Le Guillou Sandrine, Tosser-Klopp Gwenola, Le Provost Fabienne

## Abstract

In recent years, the increasing use of Next Generation Sequencing technologies to explore the genome has generated large quantities of data. For microRNAs, more and more publications have described several thousand sequences, all species included. In order to obtain a detailed description of microRNAs from the literature for three ruminant species (bovine, caprine and ovine), a new database has been created: RumimiR. To date, 2,887, 2,733 and 5,095 unique microRNAs of bovine, caprine and ovine species, respectively, have been included. In addition to the most recent reference genomic position and sequence of each microRNA, this database contains details on the animals, tissue origins and experimental conditions available from the publications. Identity with human or mouse microRNA is mentioned. The RumimiR database enables data filtering, the selection of microRNAs being based on defined criteria such as animal status or tissue origin. For ruminant studies, RumimiR supplements the widely used miRBase database by browsing and filtering using complementary criteria, and the integration of all published sequences described as novel. The principal goal of this database is to provide easy access to all ruminant microRNAs described in the literature.

## INTRODUCTION

MicroRNAs are small non-coding RNAs that are 17-22 nt in length (1). These highly conserved RNAs participate in the post-transcriptional regulation of genes through their impact on targeted messenger RNAs (mRNAs). This can lead to a repression of translation or a degradation of the targeted mRNAs depending on the binding by base-pairing of the microRNAs on their target via a recognition site, the “seed” sequence.

At present, many studies use Next Generation Sequencing (NGS) technology to explore the transcriptome, notably in a context of microRNA discovery. A significant amount of data is thus generated, which to date can reach more than a thousand microRNA sequences in a single discovery publication (2, 3). Regularly updated tools for data exploration are therefore required and essential. Animal, plant and virus microRNAs are listed in several databases, such as miRNEST 2.0 (which covers more than 400 different species) (4) or miROrtho containing 46 animal genomes (5). Some databases are more restricted to human microRNA in a context of diseases, such as EpimiRBase (microRNAs and their association with epilepsy) (6), miRCancer (microRNAs and cancer) (7) or MSDD, the MicroRNA SNP Disease Database (genetic variants affecting microRNA in a context of disease) (8). The recent miRCarta database (9) which lists the microRNAs for 148 species, is based on the prediction of novel microRNAs only. A database portal, miRToolsGallery, was recently set up and contains more than 1000 tools useful to study, identify or predict the targets of microRNAs (10). However, the most widely used database is miRBase, created in 2006 by Griffiths-Jones and collaborators at the University of Manchester (11) and listing microRNAs for 271 species (12).

Studies on bovine, caprine and ovine species have often focused on production traits such as dairy and meat products (13–15), health (mastitis resistance (16, 17)) or reproduction (fertility and fecundity (18–20)). Because of genome conservation between these three species, their miRNomes (all microRNAs expressed in a tissue or cell type) are quite similar. However there are some differences, as well as those between breeds (21). That is why it is important to generate a database on all three species which contains specific information such as the breeds of the animals or their physiological status, which are frequently lacking in the most widely employed databases.

The RumimiR database provides a single portal of access to the integration of all published information on ruminant microRNAs. RumimiR is freely available online at the following URL: http://rumimir.sigenae.org/.

This database offers an exhaustive list of bovine, caprine and ovine microRNAs, based on the data available in the literature. Moreover, pertinent information, notably in a context of dairy production, is added to the description of each mature microRNA and there is a filtering option for all these data. The entire database and data filtered in this way (using one or more filter) can be downloaded in different formats.

## MATERIALS AND METHODS

### Data collection and processing

The data included in the RumimiR database was collected from all publications describing ruminant microRNAs. Thus, data from 78 publications found in PubMed (https://www.ncbi.nlm.nih.gov/pubmed) are now present in RumimiR. The references for all these publications are listed in the Supplementary Data (**Supplementary Table 1**). Their titles and authors, with a hyperlink to the publication, are provided. RumimiR includes all known microRNA sequences, as well as those described as “novel” in the publications, which are not always found in the miRBase database. In order to standardise the data, a unique reference genome for each species is used in RumimiR. For this purpose, Blast was implemented using the NCBI tool (22), based on last versions of the reference genome available (UMD3.1.1 for bovine, ARS1 for caprine and Oar v4.0 for ovine), if this information was not mentioned in the publication, which was the case for many of them. The NCBI Blastn tool (“Somewhat similar sequences”) was used, and the general parameters applied were those by default (22). In the event of multiple alignments, with a query cover and 100% identity on the genome, a Blast of the sequence of the microRNA precursor could enable identification of its position. For 344 sequences, there were multiple positions even after applying the precursor sequence, when it was available in the publication. These microRNAs are therefore listed in the RumimiR database without their position, but the number of positions on the genome is shown in the “chromosome” column. To add information on a microRNA listed in RumimiR database, a search was made to determine its identity with a human or mouse microRNA. This identity was considered if it had a 100% sequence match with a known microRNA on the full length of the shorter sequence, using the miRBase database (release 22) (12). These identities and this step were included because of the large number of publications and hence the details available on human and mouse microRNAs. This characterisation was improved by comparing all sequences with small nucleolar RNA (snoRNA), transfer RNA (tRNA), ribosomal RNA (rRNA) and small nuclear RNA (snRNA) sequences. Indeed, part of the snoRNA structure, the stem loop, is almost identical to that of microRNA (23). tRNAs with three hairpin loops could also be confused with microRNA; to prevent this, the sequences were intersected with the list of all species snoRNA and bovine tRNA present in version 3 of snoRNABase (24) and in GtRNAdb 2.0 (25), respectively, and with the bovine, caprine and ovine rRNA and snRNA sequences present in BioMart, an Ensembl tool (26). When a microRNA displayed 100% identity with a snoRNA, tRNA, rRNA or snRNA in its full length, it was retained in the RumimiR database but this information is mentioned in the “small RNA” column. All the processes used to add microRNAs to the RumimiR database are presented in **Figure 1**.

**Figure 1.**
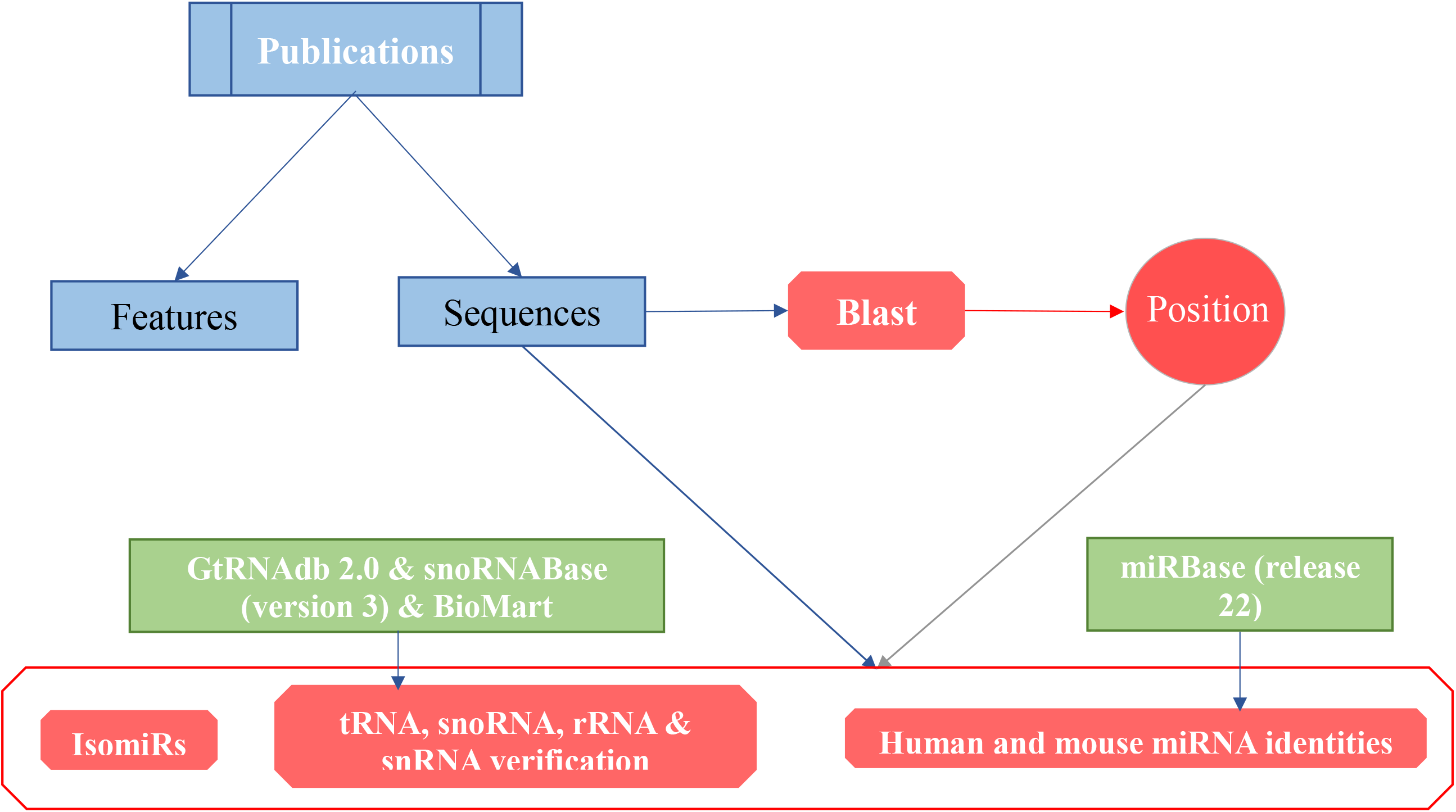
Procedure followed to include microRNAs in the RumimiR database. The features and sequences provided in publications (blue rectangles). Sequences were used to specify their genomic position from Blast analyses and to improve theirs characterisation (red forms) using the public databases available (green rectangles).

All microRNAs and most of the features described in the publications are therefore documented in the RumimiR database. All microRNA sequences are noted, as is the presence (or not) of isomiRs (microRNA sequences at the same genomic position, with almost identical sequences). In order to highlight these almost identical sequences, their names are shown in a specific column (“isomiRs”) in RumimiR. All microRNAs are therefore listed with these features and whether they belong to a microRNA family. The database also contains details of the number of animals studied, their breeds, ages and lactation stage, as well as tissues if this is mentioned in the publication.

The RumimiR website was built using HTML5 technology (https://dev.w3.org/html5), the bootstrap front-end framework (https://getbootstrap.com-v4.0.0) with additional jQuery user interface elements (http://jquery.com-v3.2.1), DataTables jQuery plug-in for the data table (https://datatables.net-v1.10.16) and the plots implemented by using the Highcharts JavaScript library (http://www.highcharts.com-v6.2.0). Rumimir has been successfully tested on Chrome (version 49 and later) and Firefox (version 57 and later). The data, in JSON format, were provided to the DataTables jQuery plug-in by setting the ajax option to the address of the JSON data source. All the statistics, the charts and drop-down lists were built on the fly based on the data to make the updates as easily as possible. The Blast tools has been developed using Perl-CGI.

## RESULTS AND DISCUSSION

### Content of the RumimiR database

At present, the RumimiR database contains 10,715 different microRNAs for the three ruminant species: 2,887 for bovines, 2,733 for caprines and 5,095 for ovines. Forty-four publications were used to collect bovine microRNAs, 19 for ovine microRNAs and 15 for caprine microRNAs (Figure 2A). The higher number of microRNAs described in ovine species was due to recent studies and hence the use of more recent NGS technologies (**Figure 2B**). More precisely, 16,041 sequences are present in the RumimiR database, corresponding to 10,715 different mature microRNAs. This difference can be explained by the presence of isomiRs. The microRNA listed in RumimiR have identities with 889 human or mouse microRNAs: 11.78% of the microRNAs described in the database present sequence identities with a human and mouse microRNA and 4.66% present sequence identity with only a human or mouse microRNA. A comparison with snoRNA, tRNA, rRNA and snRNA sequences revealed that 48 microRNAs display sequence identities with part of a tRNA, 47 with part of a snoRNA, 34 with part of a rRNA and nine with part of a snRNA. The majority of the microRNAs listed in RumimiR have the expected length (20-22 nucleotides), with 99.27% of them being 17-25 nucleotides long.

**Figure 2.**
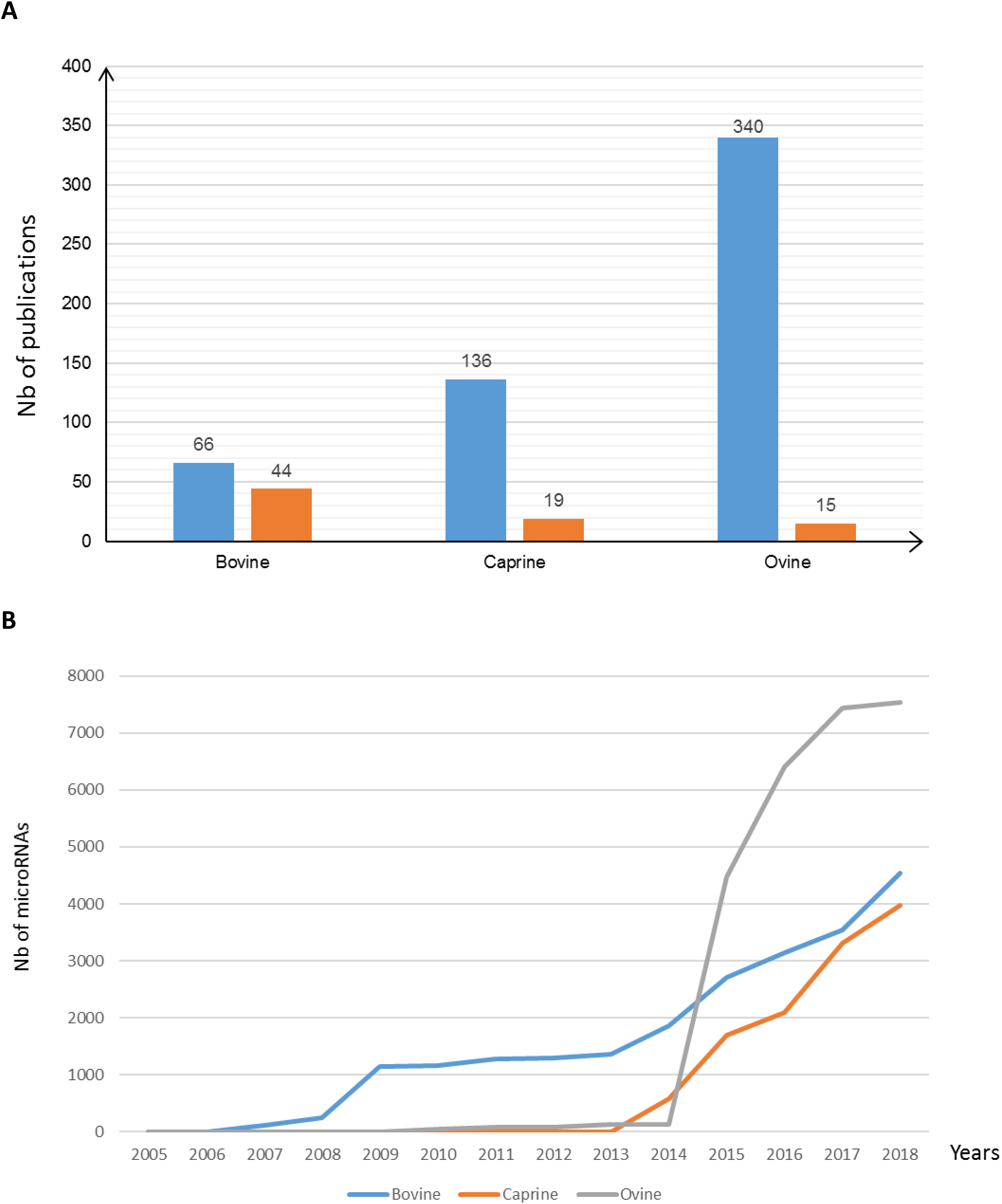
Number of microRNAs per publication. A-In blue, average number of microRNAs described per publication for each species. In orange, the number of publications on each species. B-Cumulative number of microRNAs described per year in each species.

Most of the microRNAs in the database resulted from studies on differences between breeds, developmental stages (mainly in a context of meat production) or the immune response (comparison between heathy animals and those suffering from mastitis, a pathology widely studied in ruminants) (**Figure 3**). The different issues addressed in the publications were used in to determine a filter that could be included in the RumimiR database. Forty different breeds are thus represented in the RumimiR database: 17 bovine, 14 caprine and nine ovine breeds, of both the meat and dairy breeds. About 30 tissues and body fluids are represented, such as milk (27, 28), adipose tissue (2), mammary gland (29– 31) and ovaries (20, 32, 33). More than 30 different ages are listed through their description in 38% of the publications. Some studies described the differential expression of microRNAs throughout the development and at different ages, which is why several ages might be considered in a single publication. The same was applied to lactation stages. All these figures will increase as the database is updated. The numbers of microRNAs common to the three ruminant species, or sequences common to these species and also the human and mouse, or specific of each species are presented in **Figure 4**. The percentage of specific microRNAs is between 80.24% and 88.82% for these species.

**Figure 3.**
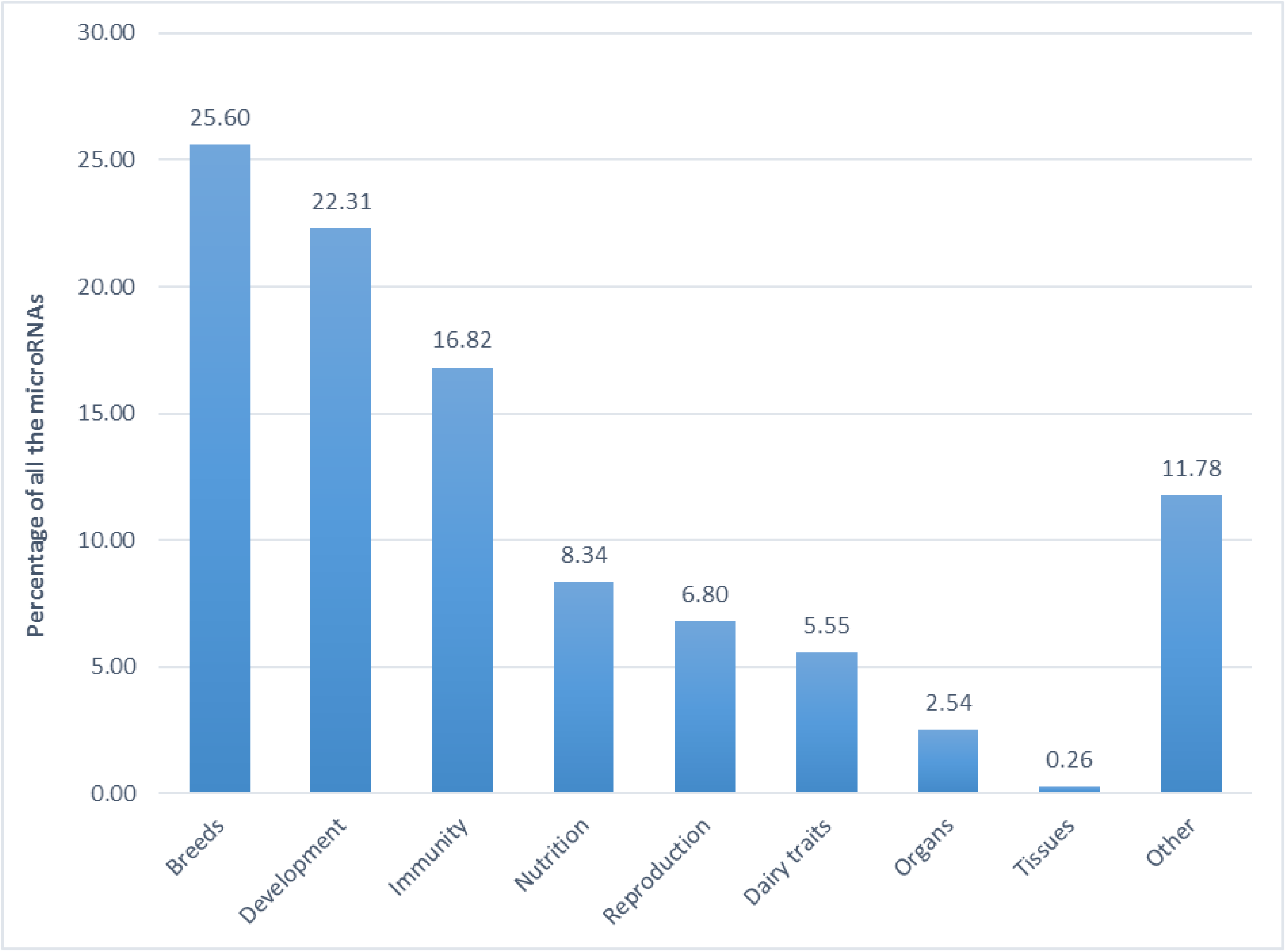
Percentage of microRNAs according to the conditions studied. The different topics presented here are in line with the issues addressed in the publications. 78 publications were considered; each article is mentioned only once.

**Figure 4.**
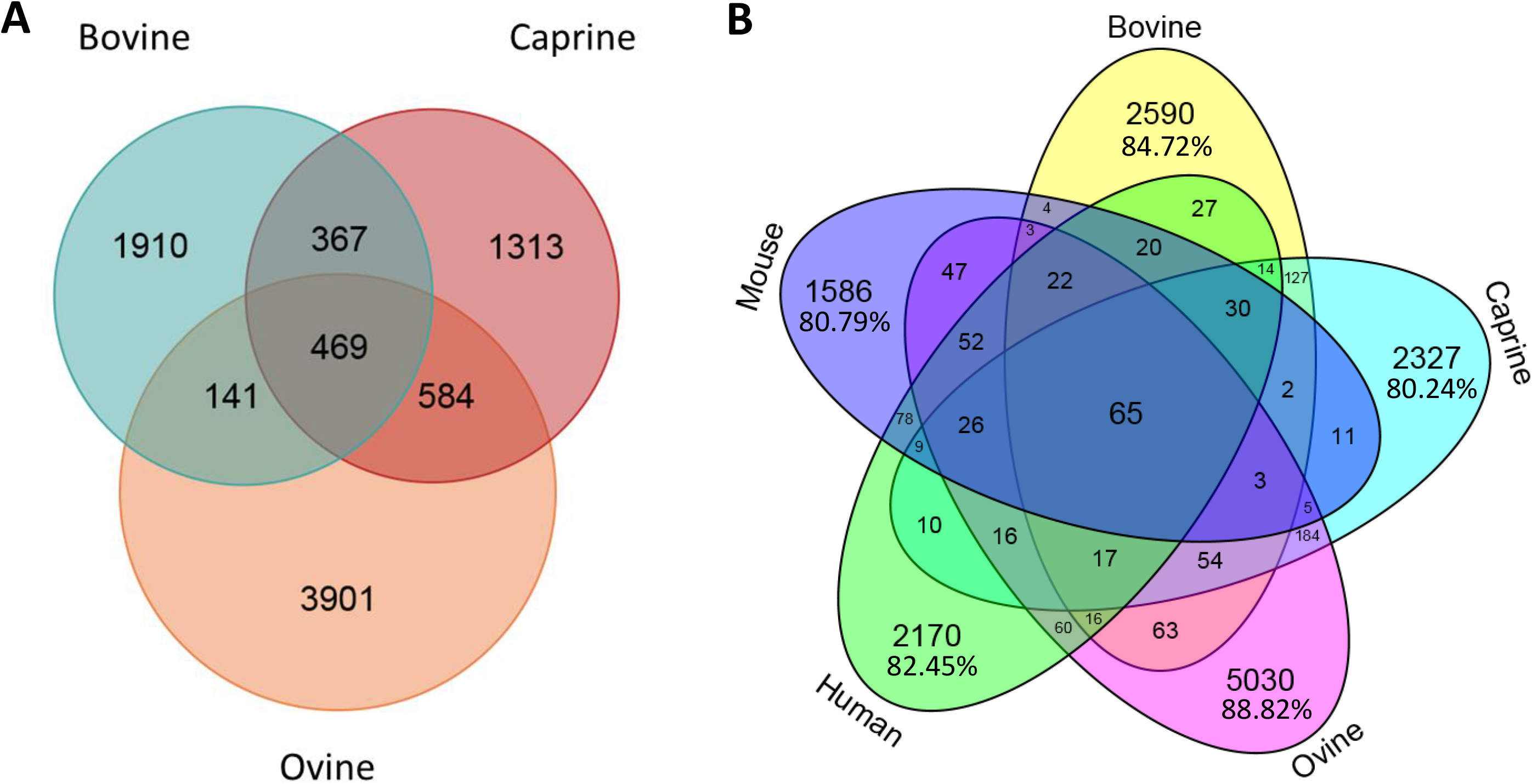
Number of microRNAs common to different species. A-Venn diagram showing bovine, caprine and ovine unique microRNAs listed in the RumimiR database. B-Venn diagram showing bovine, caprine, ovine microRNAs listed in the RumimiR database and the human and mouse microRNAs listed in miRBase (release 22).

RumimiR is limited to three ruminant species, unlike miRBase, which is the most widely used and complete database in terms of the number of species covered (plants, animals and virus). However, this restriction has enabled us to generate a more detailed database containing all the microRNAs described in different publications as well as a large number of features. The number of microRNAs listed in the RumimiR database for bovine, caprine and ovine species is six times higher than in miRBase: 10,715 in RumimiR versus 1,614 in miRBase. There remains a risk of including false positive microRNAs but RumimiR offers a more precise and complete microRNA database for these species and will therefore be of value to scientists working on livestock species.

### User interface

The web interface is user-friendly and enables visualisation of all the microRNAs described in bovine, caprine and ovine species. The position of mature microRNA (chromosome, start and end) is the minimum information provided to users. Some or all of the other columns can be selected by clicking on their name in the “show/hide columns” box (**Figure 5**).

**Figure 5.**
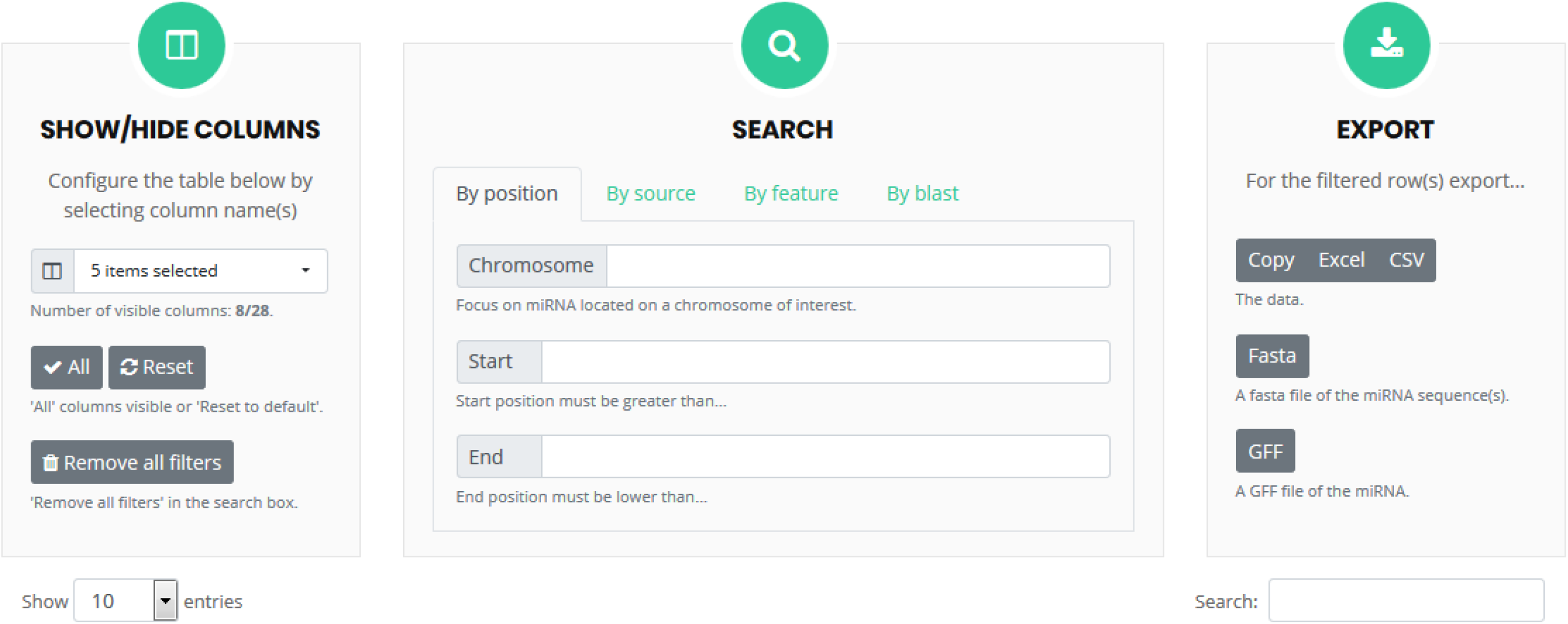
User interface for the search and browse. Columns can be selected so that only specific features will be visible. The search box enables the detection of microRNAs present in the RumimiR database as a function of the selected choices. The results can be exported in different formats, such as Excel or Fasta.

There are 28 columns which represent the features available on microRNA such as the sequence, related publications, name, isomiRs, family, tissue origin, breed, condition studied, and so on. They are all listed in **Table 1**.

**Table 1.** List of columns presenting the microRNA and features detailed in the RumimiR database. Lines in grey represent a feature present in RumimiR but not in miRBase.

|  |  |
| --- | --- |
| Chr | Genomic position of the microRNA |
| Start |  |
| End |  |
| Sequence | Sequence of the mature microRNA |
| Publication | Publication in which the microRNA is described |
| Name | Name assigned to the microRNA |
| IsomiRs | Name of isomiRs of the microRNA (sequence and genomic position almost identical) |
| Family | microRNA family to which the microRNA is affiliated |
| Tissue | Tissue in which the microRNA was discovered |
| Breed | Breed used during the study |
| Study conditions | Conditions prevailing during the study |
| Lactation stages | Lactation stages of the animals studied |
| Age | Age of the animals studied |
| Species | Species of the animals studied (bovine, caprine or ovine) |
| Number | Number of novel microRNAs described in the publication |
| Cut-off | Cut-off point used in the publication to detect novel microRNAs |
| Method | Method used to detect novel microRNAs |
| Number and breed | Number of animals studied in each breed |
| Reference genome | UMD3.1.1, ARS1 or Oar v4.0 |
| Bioinformatics tools used | Software used in the study for the microRNA detection |
| Condition details | Details of study conditions (i.e. number of animals per condition) |
| 5p/3p | If the microRNA is 5p or 3p |
| Strand (+/-) | Strand in which the microRNA is situated |
| Star sequence | Sequence of the star microRNA |
| Matching seed | Known microRNA with the same seed |
| small RNAs | If the microRNA described is in fact part of a snoRNA, tRNA, snRNA or rRNA |
| hsa homology | Similarity with a human microRNA |
| mmu homology | Similarity with a mouse microRNA |

Users can also visualise just those microRNAs that are of interest to them by selecting the column corresponding to their request: the sequence and/or tissue and/or breed, etc., and then using the “search” box to define different options such as “by position” (chromosome, start, end), “by source” (species, breed, tissue) or “by feature” (conditions, method, software) (**Figure 5**). It is also possible to filter all the mature microRNAs listed, according to the different criteria mentioned above. A sequence or key word can also be entered in the “Search” box, and the relevant data will be listed in the results table. When a selection of microRNAs with the requested specificities is available, the data thus filtered are apparent and downloadable in several formats (Excel, CSV, Fasta or GFF). Users can thus rapidly obtain all the data required, appropriately filtered. It is also possible to download the entire database without any filters being applied.

For improved visibility, certain statistics and graphs are presented in the online version, summarising the data presented in the database. The date of the last update is mentioned, as is the number of sequences listed and the number of publications used, amongst other items. The distribution of microRNA by species and by breed is presented graphically, as is the distribution by tissue origin.

An alignment tool is also available in the RumimiR database; sequences can be submitted and the microRNAs corresponding to these sequences are then sorted by hits.

### Future extensions

The NGS approach generates extensive data and the number of microRNAs will continue to rise in the near future. We will continue to integrate new data and update the database on a regular basis so that it remains exhaustive. One potential extension for RumimiR might be the addition of genetic variants of microRNAs in ruminant species (“miRSNPs” for microRNA and Single Nucleotide Polymorphisms); firstly those linked with dairy QTL, and secondly with health or meat QTL. Publications on these genetic microRNAs variations are indeed increasingly numerous. For example, in humans, the MSDD database has been created exclusively for miRSNPs linked to human diseases (8). In the same way, the integration of ruminant miRSNPs will enable completion of the RumimiR database. Other features and filters may also be added in accordance with the features found in publications and the needs of the scientific community. The evolution of reference genomes will also be considered, to take account of the genomic position microRNAs in the latest versions of the reference genomes. The RumimiR database could also be extended to include other livestock species.

## Conclusion

The RumimiR database contains an exhaustive list all of the microRNAs described in the literature for three livestock species: bovine, caprine and ovine species. This database supplements one of the most widely used databases on microRNAs, miRBase, thanks to the availability of various features mentioned in the publications which are of importance in the context of animal production and dairy traits, notably the breeds of the animals studied or the origins of tissues in which the microRNAs have been described. RumimiR enables the retrieval of specific microRNAs and thus provides additional details on livestock species that are necessary to gaining a clearer understanding of the context in which they were discovered. The database will be regularly updated and thus continue to be exhaustive. RumimiR offers a unique tool that presents and describes all the microRNAs discovered by standardising and centralising information from a large number of publications.

## FUNDING

This work was supported by APIS-GENE through the miRQTLait project. CB’s grant was financially supported by INRA and APIS-GENE.

## ACKNOWLEDGEMENTS

The authors are grateful to C. Gaspin, C. Leroux and S. Marthey for their critical reading of the manuscript and to M. Boussaha for his helpful discussions.

## SUPPLEMENTARY DATA

Supplementary Table 1. List of publications used to collect microRNA data.

For this first version of RumimiR database, the microRNAs from 78 publications were collected and included in the database. Publications 1 to 44 correspond to bovine studies, 45 to 63 to caprine studies and 64 to 78 to ovine studies.

