## Supplemental Table 1 for "RumimiR: a detailed microRNA database focused on ruminant species"

27. Malvisi,M., Palazzo,F., Morandi,N., Lazzari,B., Williams,J.L., Pagnacco,G. and Minozzi,G. (2016) Responses of Bovine Innate Immunity to Mycobacterium avium subsp. paratuberculosis

Infection Revealed by Changes in Gene Expression and Levels of MicroRNA. *PLOS ONE*, **11**, e0164461.
